# NCBP3 is a productive mRNP component

**DOI:** 10.1101/2020.07.04.184861

**Authors:** Yuhui Dou, Isabelle Barbosa, Hua Jiang, Claudia Iasillo, Kelly R. Molloy, Wiebke Manuela Schulze, Stephen Cusack, Manfred Schmid, Hervé Le Hir, John LaCava, Torben Heick Jensen

**Affiliations:** Department of Molecular Biology and Genetics, Aarhus University, C.F. Møllers Allé 3, Building 1130, Aarhus 8000, Denmark; Institut de Biologie de l’ENS (IBENS), Département de biologie, École normale supérieure, CNRS, INSERM, Université PSL, 75005 Paris, France; Laboratory of Cellular and Structural Biology, The Rockefeller University, 1230 York Avenue, New York, NY 10065, USA; Laboratory of Mass Spectrometry and Gaseous Ion Chemistry, Rockefeller University, 1230 York Avenue, New York, NY 10065, USA; European Molecular Biology Laboratory, Grenoble Outstation, 71 Avenue des Martyrs, CS 90181, Grenoble Cedex 9 38042, France; European Research Institute for the Biology of Ageing, University Medical Center Groningen, Groningen 9713 AV, Netherlands

## Abstract

The nuclear Cap Binding Complex (CBC), consisting of Nuclear Cap Binding Protein 1 (NCBP1) and 2 (NCBP2), associates with the nascent 5’cap of RNA polymerase II transcripts and impacts RNA fate decisions. Recently, the C17orf85 protein, also called NCBP3, was suggested to form an alternative CBC by replacing NCBP2. However, applying protein-protein interaction screening of NCBP1, 2 and 3, we find that the interaction profile of NCBP3 is distinct. Whereas NCBP1 and 2 identify known CBC interactors, NCBP3 primarily interacts with components of the Exon Junction Complex (EJC) and the TRanscription and EXport (TREX) complex. NCBP3-EJC association *in vitro* and *in vivo* requires EJC core integrity and the *in vivo* RNA binding profiles of EJC and NCBP3 overlap. We further show that NCBP3 competes with the RNA degradation factor ZC3H18 for binding CBC-bound transcripts, and that NCBP3 positively impacts the nuclear export of polyadenylated RNAs and the expression of large multi-exonic transcripts. Collectively, our results place NCBP3 with the EJC and TREX complexes in supporting the productive fate of mRNA.

## Introduction

All eukaryotic RNA polymerase II (RNAPII) transcripts undergo processing and protein-RNA packaging events, that are essential for their downstream cellular fates (1). Shortly after initiation of RNAPII transcription, a 7-methylganosine (m_7_G) cap is enzymatically added to the emerging 5’end of the nascent RNA. While providing protection against 5’-3’ exonucleolysis, the cap also serves as a binding site for the Cap Binding Complex (CBC), which is composed of the Nuclear Cap Binding Proteins 1 and 2 (NCBP1/CBP80 and NCBP2/CBP20, respectively) and considered a hallmark of all nuclear RNAPII transcripts (2). NCBP2 directly contacts the 5’cap, through its RNA Recognition Motif (RRM), however, high affinity cap-binding is only achieved upon dimerization of NCBP2 and NCBP1 (3,4). Attached to NCBP2, the larger NCBP1 protein serves as a platform for the sequential formation of different complexes engaged with ribonucleoprotein particle (RNP) biology (3,5). In this sense, the CBC impacts various aspects of gene expression, from transcription (6), RNA splicing (2), RNA 3’end processing (7,8), RNA transport (9,10) to RNA decay (11–13), and is hereby capable of influencing RNP fate in both productive and destructive ways (14).

The impact of the CBC on RNP identity is diverse. For example, early in the RNP assembly process, the CBC may interact with the ARS2/SRRT protein to form the CBC-ARS2 (CBCA) complex and further with PHAX, forming the CBC-ARS2-PHAX (CBCAP) complex (12,15,16). If the transcription unit (TU) produces an snRNA, or an independently transcribed snoRNA, transcription termination-coupled RNA 3’end processing will, together with CBC-bound PHAX, promote the nuclear export or intranuclear transport of the resulting snRNPs and snoRNPs, respectively (9,17). If the TU produces an intron-containing RNA, the CBC will positively impact splicing by interacting with the U4/U6.U5 tri-snRNP and promote spliceosome assembly (2,18). The spliced RNA will then be marked by the Exon Junction Complex (EJC), composed of its four core components EIF4A3, RBM8A/Y14, MAGOH and MLN51/CASC3/BTZ, as well as its multiple peripheral factors (19,20). In addition to the EJC, the Transcription and EXport (TREX) complex is recruited co-transcriptionally (21,22). A central TREX component ALYREF can be recruited by the CBC directly to the 5’ end of RNA, and joined by other TREX proteins, e.g. DDX39B and the THO complex, together with the EJC, leading to the formation of a nuclear export competent mRNP (10,22–24).

In addition to its effects on RNP production and function, the CBC may also promote RNA nuclear degradation. In the case of such short-lived transcripts, the CBCA complex can link to the ZC3H18 protein and further to the Nuclear EXosome Targeting (NEXT) complex (12,25) or the PolyA eXosome Targeting (PAXT) connection (26), which both promote RNA degradation via the nuclear RNA exosome (27). Given this broad repertoire of CBC interactions, it is a central question in nuclear RNA biology how RNP identity is ultimately established. Here, it has been suggested that nuclear RNAs are constantly targeted by factors, often involving mutually exclusive CBC-containing complexes, which ultimately ‘settle’ RNP fate commitment as a consequence of still ill-defined molecular decision processes (28). Such competing interactions are for example illustrated by the competition between the NEXT/PAXT component MTR4 and ALYREF for association with CBCA-bound RNAs (29) and by the mutually exclusive CBC interactions of ZC3H18 and PHAX (13).

In addition to NCBP1 and NCBP2, the C17orf85/ELG protein was recently shown to bind the m_7_G cap through its putative N-terminal RRM (30). It was further proposed to be able to substitute for NCBP2 in forming an alternative CBC by interacting with NCBP1 via its C-terminal region (30,31); hence it was renamed NCBP3. However, the cap-affinity of NCBP3, unlike that of NCBP2, was not enhanced by NCBP1, and the NCBP1-NCBP3 complex bound the m_7_GTP cap analogue with ~50 fold weaker affinity than that of the canonical CBC (32). Moreover, NCBP3’s association with NCBP1 was not mutually exclusive, but rather compatible, with that of NCBP2 (30,32). Consistent with NCBP3 being a cellular partner of ARS2 (30,32), the protein was capable of interacting with the CBCA complex *in vitro* and in a manner preventing formation of the CBCAP complex (32). Considering the central concept of factor competition in determining nuclear RNP identity, and to interrogate the alternative CBC hypothesis, we therefore aimed to investigate NCBP3 interactions further.

Applying extensive immunoprecipitation (IP) screening of NCBP1, NCBP2 and NCBP3, we demonstrate a distinct protein interaction network of NCBP3 as compared to its canonical CBC counterparts. Our results identify components of the EJC and TREX complexes as abundant NCBP3 interactors and we show that NCBP3 accumulates in nuclear speckles together with these components. Interestingly, NCBP3 associates with the assembled EJC core and binds RNAs in proximity to the EJC. NCBP3 further aids the TREX complex in mRNA export and positively regulates the expression of multi-exonic RNAs. We conclude that NCBP3 helps to promote the productive fate of mRNPs, which is supported by its ability to compete with ZC3H18 for RNP association.

## Materials and Methods

### Cell lines and cultures

HeLa Kyoto cells stably expressing LAP-tagged proteins (NCBP1, NCBP2, NCBP3 and ALYREF) and control LAP cells were kindly provided by Anthony Hyman (33) (see **Supplementary Table 4**). NCBP1-, NCBP2-, NCBP3- and control-LAP cells were FACS sorted for EGFP-positive cells. HeLa cells expressing endogenously HA-tagged EIF4A3 were from (51). All cells were maintained in Dulbecco’s modified eagle medium (Gibco) supplemented with 10 % fetal bovine serum (Sigma) and 100 U/ml penicillin and 100 μg/ml streptomycin (Sigma), at 37 °C and 5 % CO_2_.

### RNA interference

RNA interference (RNAi) was performed with Lipofectamine 2000 (Invitrogen) according to the manufacturers’ instructions. The sequences of the used siRNAs are shown in **Supplementary Table 5**. Cells were seeded 16 h prior to transfection with Lipofectamine 2000 (10 μl/ 10 cm dish) and siRNAs (final concentration of 20 nM). 48 h later the transfection was repeated. Cells were generally harvested 24 h after the second transfection, but for experiments with depletion of EIF4A3, RBM8A, ALYREF and DDX39B, cells were harvested 48 h after the first transfection. For cells used for the RNA-seq analysis, 15 nM of siRNAs was used and cells were harvested 36 h after the second transfection. Depletion efficiencies were monitored by western blotting analysis and protein extractions were performed as described in ‘Co-IP experiments’ below.

### Co-IP experiments

The interaction screening was performed essentially as described in (34), (see Dou et al., submitted, for more details). Briefly, 3 or 4 replicate IPs were conducted, for each of the extraction conditions. IPs of LAP-tagged NCBP1, NCBP2 and NCBP3 cell lines were carried out in parallel with IP using CTRL-LAP cells. Protein extractions were performed differently for the interaction screen IPs (a) and for the individual IPs (b). (a) Proteins were extracted from cryogrinded cell powders with different extraction solutions supplemented with protease inhibitors (Roche) (solution compositions listed in **Supplementary Table 1**), at 1:9 (w/v) for 50 mg of NCBP1 and NCBP2, and 1:4 (w/v) for 300 mg of NCBP3, with the aid of brief sonication (QSonica Q700, 1 Amp, 30~40 s; or QSonica S4000, 2 Amp, 15 x 2 s). (b) Cells were lysed in extraction solution (20 mM HEPES, pH 7.4, 0.5 % TritonX-100, 150 mM NaCl, supplemented with 1x protease inhibitors), and sonicated with a microtip sonicator (Branson 250) for 3 x 3 s. After sonication, for both interaction screen IPs and individual IPs, extracts were clarified by centrifugation at 14,000 g and 4 °C for 10 min. Supernatants were then incubated with anti-GFP antibodies conjugated to Dynabeads Epoxy M270 (Invitrogen) (69) and rotated at 4 °C for 30 min; or anti-HA antibodies (Abcam, #ab9110) conjugated to protein A Dynabeads (Invitrogen) for EIF4A3-HA IPs and rotated at 4°C for 2 h. Beads were subsequently washed three times with extraction solution and captured proteins were eluted with 1.1 x LDS for SDS-PAGE.

Where RNase treatment was applied, the beads were, after three times of washing, resuspended in 40 μl extraction solution with 1 μl of RNase A/T1 mix (Thermo Scientific # EN0551), and left shaking for 20 min at room temperature, and washed once before elution.

### SDS-PAGE and western blotting analysis

NuPAGE 4-12 % Bis-Tris gels (Invitrogen) were run with MOPS or MES buffers (Invitrogen), depending on the sizes of the target proteins, and following the manufacturer’s manual. Proteins were subsequently transferred to a 0.45 μm PVDF membranes (Immobilon-P membrane) with the iBlot2 transfer system (Thermo Fisher), and blocked with 5% BSA in Tris-buffered saline and 0.1% Tween 20 (TBST). Membranes were then incubated with primary antibody (in TBST with 1% BSA) (antibody list in **Supplementary Table 6**) at 4°C overnight, followed by washing 3 x 10 min with TBST. HRP-conjugated anti-mouse or rabbit secondary antibodies (Dako), or HRP-conjugated VeriBlot IP Detection Reagent (Abcam) were then applied (1: 10000 in TBST with 1 % BSA), and incubated at room temperature for 1 h. After washing as above, membranes were incubated with ECL substrate (Thermo Scientific #34096) and imaged with Amersham Imager 600.

### MS and data processing

Details of MS sample processing is described elsewhere (Dou et al., submitted). Briefly, samples were reduced, alkylated and in-gel trypsin digested. Peptides were then extracted with 1 % TFA and cleaned with C18 tips, after which they were analysed with liquid chromatography and MS on Orbitrap Fusion or Q Exactive instruments (Thermo Fisher Scientific).

MS raw data were processed with the MaxQuant package, with “label-free quantification (LFQ)” and “match between runs” enabled (70). The Maxquant output was processed with Perseus software, where proteins labelled as “only identified by site”, “reverse”, or “contaminant” were filtered out. LFQ intensities were log2 transformed and proteins were filtered with valid values in at least two out of all of the replicates. The missing values were imputed from normal distributions with default Perseus setting. T tests were then performed between each NCBP IP and the control IP. Proteins with a false discovery rate (FDR) < 0.01, S_0_ (Log2 fold changes) > 1 were considered as significantly enriched. Using significantly enriched proteins, stoichiometric abundances were calculated as follows (36): 1) Mean LFQ intensity of the replicates was divided by the molecular weight of the protein; 2) the background of the control IP was subtracted; 3) the resulting value was normalised to that of the bait, which was set to 100. Hierarchical clustering was performed using log2 (stoichiometric abundance x 10_3_) with Euclidean distances under Perseus default settings. The Venn diagram is generated with the BioVenn tool (71).

### Recombinant proteins binding assay

The purified recombinant proteins used for interaction assays were previously described (32,40). Interaction assays were mainly performed as previously described (40). Briefly, 3 μg of each protein were mixed in Binding Buffer BB-125 in a final volume of 60 μl (125 mM NaCl final concentration) complemented, or not, with 2 mM ADPNP and/or single-stranded biotinylated RNA (10 μM). After incubation for 20 min at 30 °C, 12 μl of pre-blocked m7G affinity beads (50 % slurry, 7-Methyl-GTP Sepharose^®^ 4B, GE Healthcare) and 200 μl of BB-250 (250 mM NaCl) were added. After gentle rotation for 2 h at 4 °C, the beads were washed three times with BB-150 (150 mM NaCl) and eluted with 1X SDS loading dye. Eluates were boiled and loaded on 4-12 % SDS-PAGE with a protein marker (PageRuler unstained Protein Ladder, Fermentas). Proteins were visualized by Coomassie staining.

### Immunofluorescence and microscopy

For protein immunofluorescence, cells were grown on 18 mm cover slips, washed twice with PBS and fixed with 4 % paraformaldehyde in PBS for 12 min. They were then washed twice with PBS, permeabilized and blocked with 0.5 % Triton X-100 and 3 % FBS in PBS for 15 min and washed three times with PBS. Cells were then incubated with primary antibodies for 1 h and washed three times with PBS before incubation with secondary antibodies for 45 min followed by three times of washing with PBS (antibody list shown in **Supplementary Table 7**). Finally, cells were stained with DAPI 1 μg/ml for 10 min, followed by washing with PBS twice and H_2_O once before mounting on slides with ProLong Gold antifade reagent. PA_+_ RNA fluorescent in situ hybridization (FISH) analysis was performed as previously described (65) using a Cy5 labelled oligo (dT)70 probe (DNA Technology A/S). Images were obtained with a ZEISS Axio Observer 7 microscope equipped with an Axiocam 702 camera and Plan-APOCHROMAT 63 x/1.4 oil objective, and Zen 2 software.

### RNA extraction and RT-qPCR analysis

For RNA used for RT-qPCR analysis, RNA extraction was performed using TRIzol (Invitrogen) following the manufacturer’s instructions. RNA was DNase treated with TURBO DNA-free kit (Invitrogen) before reverse transcription with SuperScript III Reverse Transcriptase (Invitrogen). The cDNA obtained were used for qPCR with SYBR Green qPCR SuperMix (Invitrogen) and AriaMx Real-time qPCR System (Agilent). Sequence of primers used is listed in **Supplementary Table 8**.

### RNA sequencing (RNA-seq) and data analysis

RNA extraction and RNA-seq library preparation upon GFP and NCBP3 depletion were performed as described (53). Total RNAseq of siNCBP3 depletion samples are first reported here, but were collected as part of same experiment described in (53,54). Reads from siNCBP3 samples were processed in parallel with the siEGFP control and relevant depletions from GEO:GSE99059 were processed in parallel as described in (72). In brief, raw reads were quality filtered and trimmed as described (26), using Trimmomatic (v 0.32) and settings (PEILLUMINACLIP:/com/extra/Trimmomatic/0.32/adapters/TruSeq3-PE-2.fa:2:30:10 HEADCROP:12 LEADING:22 SLIDINGWINDOW:4:22 MINLEN:25). Cleaned reads were then mapped to GRCh38 with HISAT2 (v 2.1.0) (73) using default settings and the genome index ‘H. sapiens, UCSC hg38 and Refseq gene annotations’ provided at the HISAT2 download page (ftp://ftp.ccb.jhu.edu/pub/infphilo/hisat2/data/hg38_tran.tar.gz). Only proper pairs with both reads mapping to the genome were used for further analysis. Exon read counts were collected using the featureCounts tools from the subread software suite (v 2.0.0) (74) with settings ‘-p -C -s 2 -t exon’ for the default GRCh38 annotations for featureCounts. Differentially expressed transcripts were obtained from raw read counts using R package DESeq2 (v 1.20.0) at a false discovery rate (FDR) cutoff of 0.1. Exon counts for differentially up- and down-regulated transcripts were compared using custom scripts in R.

### CLIP data analysis

Processed CLIP data for NCBP3 and EIF4A3 were obtained from (46). Data shown are from NCBP3 PAR-CLIP data processed with the PARalyzer tool and the EIF4A3 HITS-CLIP data was processed with Piranha (46). NCBP2 CLIP data was from GSE94427 (13) and lifted to GRCh38. ALYREF CLIP data was from GSE113896 (22) and lifted to GRCh38.

Metagene plots for CLIP data were constructed using custom python and R scripts employing deeptools software (v3.0.2) (75) relative to GRCh38 annotations shipped with featureCounts used for RNAseq analysis.

For T>C conversion positions, raw reads were obtained from SRA: SRR500480 and SRR500481 (47), adapter sequences (TCTTTTATCGTATGCCGTCTTCTGCTTG and TCTCCCATCGTATGCCGTCTTCTGCTTG), low quality and short reads removed using bbduk from the BBMAP software (v35.92, https://jgi.doe.gov/data-and-tools/bbtools/) using parameters *k=13; ktrim=r; useshortkmers=t; mink=5; qtrim=t; trimq=10; minlength=20; ref=barcodes.fa*, where the ‘barcodes.fa’ file contains the 2 adapter sequences. Remaining reads were mapped to GRCh38 using the STAR aligner (v2.5.2b) (76) together with samtools (v1.6) (77) with settings --outFilterType BySJout; --outFilterMultimapNmax 20. Duplicate reads were removed using samtools rmdup, remaining mismatched bases collected using samtools mpileup and T to C conversion positions extracted from mpileup output using a custom script counting T>C mismatches for plus strand conversions and A>G mismatches for minus strand conversions.

## Results

### Interrogating interactions facilitated by NCBP1, NCBP2 and NCBP3

If NCBP1 and NCBP3 would form an alternative CBC (30), we surmised that their interaction would be independent of, or perhaps even competitive with, NCBP2. We therefore conducted co-IP experiments of NCBP1, employing HeLa cells that express Localization and Affinity Purification (LAP)-tagged NCBP1, upon siRNA-mediated individual depletion of NCBP2 or NCBP3. The NCBP1-NCBP3 interaction was virtually lost upon NCBP2 depletion (Fig. 1a, compare lanes 6 and 7). Moreover, the NCBP1-NCBP2 interaction was unaffected by NCBP3 depletion. Thus, despite *in vitro* studies showing that NCBP3 can bind NCBP1 in the absence of NCBP2 (30,32), it appears that the NCBP1-NCBP3 interaction *in vivo* requires the presence of NCBP2.

**Fig. 1.**
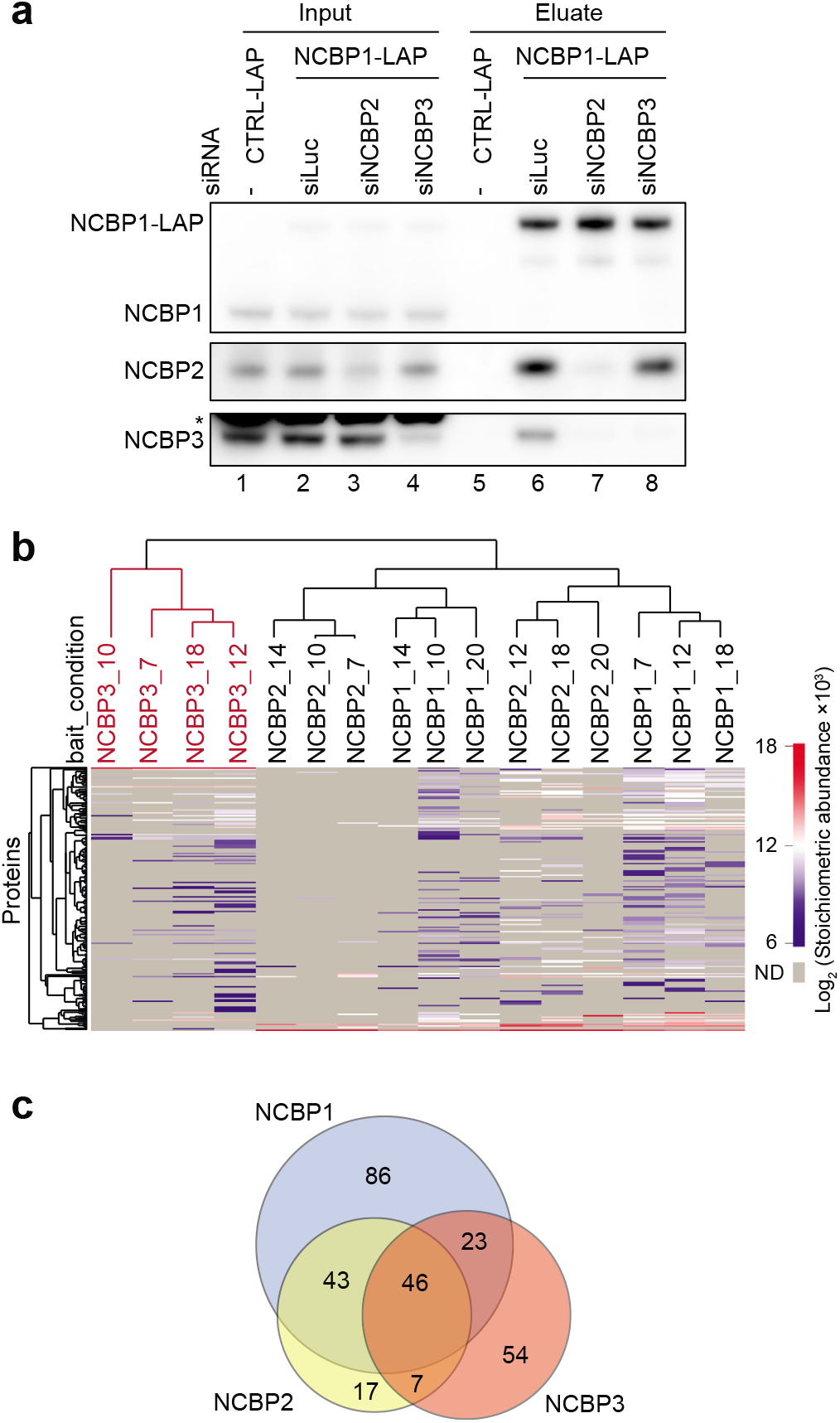
Protein-protein interactions facilitated by NCBP1, NCBP2 and NCBP3. (A) Western blotting analysis of NCBP1-LAP co-IPs from HeLa cell extracts upon depletion of luciferase (siLuc) control, NCBP2 or NCBP3 using the indicated siRNAs. IPs were performed with anti-GFP antibodies. A negative control IP (‘CTRL-LAP’) was conducted using cells expressing the untagged LAP moiety. Lanes 1-4 and lanes 5-8 show input and eluate samples, respectively. Asterisk indicates unspecific protein band. Gel loading: 0.05 % of input, 15 % of eluate. (B) Hierarchical clustering (Euclidean distances) of log2 transformed stoichiometric abundances derived from NCBP1, NCBP2 and NCBP3 interaction screening over various extraction conditions (see Supplementary Table 1). Columns represent the clustering of individual NCBP IPs (with each extraction condition signified by a number) with the NCBP3 cluster highlighted in red. Rows indicate the protein interactors identified. To calculate stoichiometric abundance, mean Label Free Quantification (LFQ) intensity was normalized to the molecular weight of the protein and deducted that of the negative control sample, the value of which was then further normalized to that of the relevant bait protein. Only proteins passing a T-test significance threshold of (FDR< 0.01 and log2 fold change> 1) were considered. (C) Venn diagram displaying the overlap of proteins captured by NCBP1-LAP (blue), NCBP2-LAP (yellow) and NCBP3-LAP (red) that were significantly enriched in at least one of the 6 extraction conditions.

As we wanted to further explore the interaction networks of NCBP1, NCBP2 and NCBP3, we next conducted IP - Mass Spectrometry (IP-MS) analysis using HeLa cells stably expressing LAP-tagged versions of the three NCBPs at near endogenous level (Supplementary Fig. 1), or the LAP tag alone as a negative control (33). To obtain comprehensive interaction profiles, we applied an optimized IP screen, where multiple extraction conditions were used to favour a broad range of protein interactions (34) Details of this IP screen are described elsewhere (Dou et al., submitted). After a preliminary screen with 24 different extraction conditions, final IPs were conducted in triplicate or quadruplicate using 6 different conditions for NCBP1 and NCBP2, and 4 different conditions for NCBP3 and the control IP (see Supplementary Table 1 for solution details). Proteins that were enriched with statistical significance in NCBP-LAP IPs over the LAP control IPs, were identified (35). Their stoichiometric abundances, relative to the relevant bait protein, were then calculated (36) and used to perform hierarchical clustering over the different conditions. Remarkably, the NCBP3 interaction profile formed a distinct cluster, whereas NCBP1 and NCBP2 interactions clustered together (Fig. 1b). Consistently, ~80% of proteins captured by NCBP2 were common with NCBP1, whereas NCBP3 only shared ~50% of its interactors with NCBP1, with the majority of the rest being NCBP3-specific (Fig. 1c).

Taken together, our results demonstrate that NCBP3 harbours a partially distinct interaction profile and suggest that the protein only associates with NCBP1 and NCBP2 in their CBC context.

### NCBP3 predominantly interacts with EJC and TREX components

Further analyses of the stoichiometric interaction profiles of NCBP proteins revealed that NCBP1 and NCBP2 primarily interacted with each other and with PHAX (Supplementary Fig. 2). In contrast, the most predominant interactors of NCBP3 were components of the EJC and TREX complexes, among which EJC core components EIF4A3, RBM8A and MAGOH were most abundantly enriched across all IP conditions (Supplementary Fig. 2). To better compare interaction differences between each of the NCBP proteins, we summarized protein groups/complexes that abundantly interact with NCBP3 or NCBP1 and NCBP2 in a heatmap (Fig. 2a, for extensive complex enrichment comparison, see Dou et al., submitted). PHAX was exclusively, and ARS2 was primarily, enriched in NCBP1 and NCBP2, but not in NCBP3, IPs, which was consistent with previous studies (30,32). Another group of proteins that stood out as being enriched in NCBP1 and NCBP2, but not in NCBP3, IPs, were factors related to RNA decay by the nuclear exosome, including ZC3H18, MTR4/SKIV2L2 and ZCCHC8. The heatmap displays that EJC and TREX components are abundant NCBP3 interactors exhibiting substantially enhanced co-enrichment with NCBP3 compared to NCBP1 and 2.

**Fig. 2.**
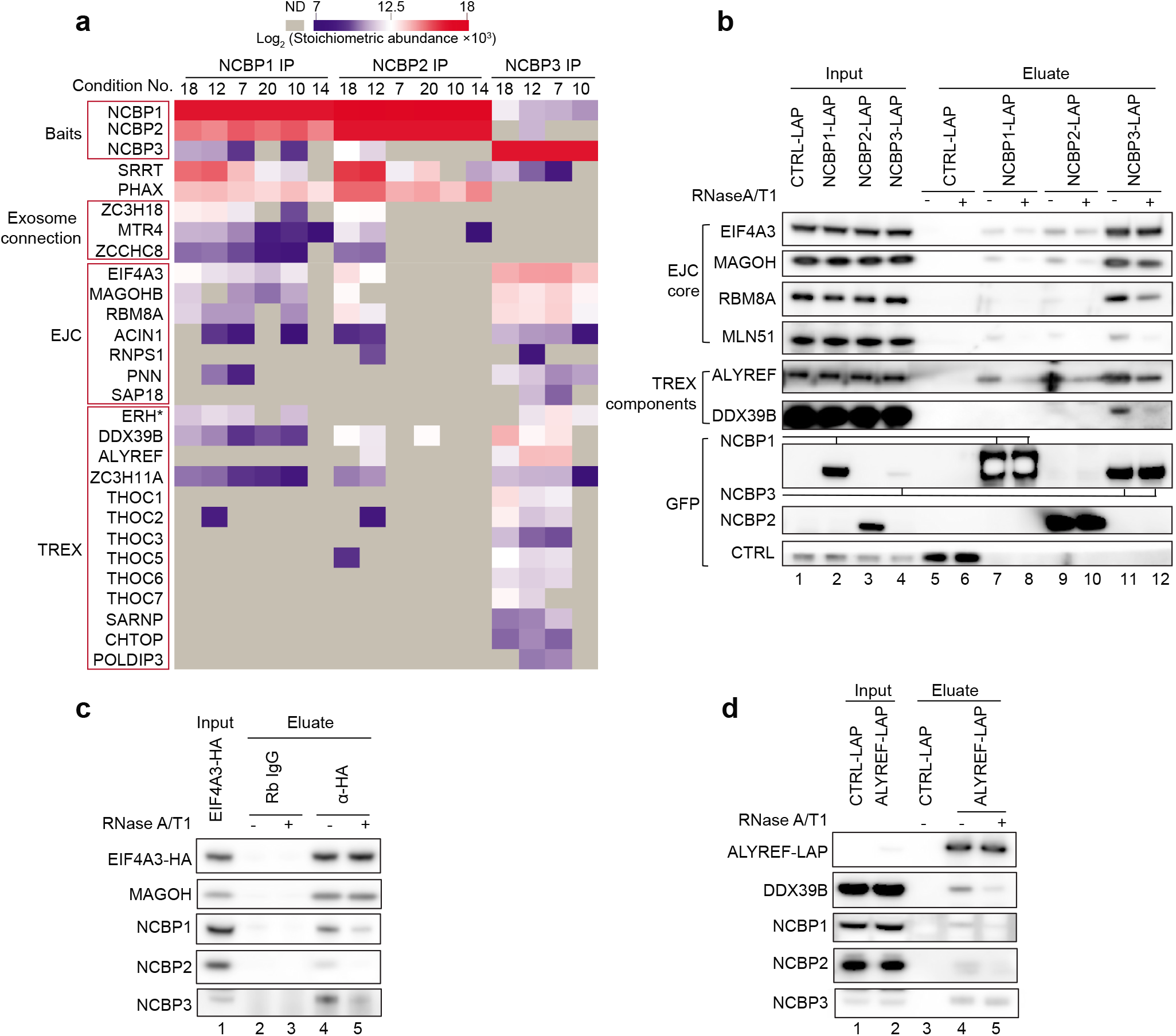
NCBP3 predominantly interacts with EJC and TREX components. (A) Heatmap comparing abundances of selected protein groups among the three NCBP protein IPs as indicated. The heatmap shows log2 transformed stoichiometric abundances of proteins significantly enriched in the NCBP IPs across different extraction conditions. Proteins known to belong to certain complexes are grouped in labelled boxes on the left. The heat colour range is shown on the top and missing values (ND) are shown in grey. *ERH as a putative TREX components (21,41). The full data set of significantly enriched proteins is provided in **Supplementary Table 2**. (B) IP/western blotting analysis validating the interactions of NCBP proteins with the indicated EJC and TREX components. IPs were performed on extracts from control (CTRL-LAP), NCBP1-LAP, NCBP2-LAP and NCBP3-LAP HeLa cells with (+) or without (-) RNase A/T1 treatment. Bait proteins were detected with anti-GFP antibodies. Lanes 1-4 and 5-12 show input and eluate samples, respectively. Gel loading: 0.05% of input, 15% of eluate. (C) Western blotting analysis of the indicated proteins after EIF4A3-HA co-IP with HA antibody (Rabbit IgG as negative control). RNase A/T1 treatments were as indicated. Gel loading: 0.05% of input, 15% of eluate. (D) Western blotting analysis of the indicated proteins after ALYREF-LAP co-IP with anti-GFP antibody (CTRL-LAP cell extract as negative control). RNase A/T1 treatments were indicated. Gel loading: 0.05% of input, 15% of eluate.

We next sought to confirm the interactions of NCBP3 with EJC and TREX components by IP/western blotting analysis. While all three NCBP proteins were able to co-IP EIF4A3, RBM8A and MAGOH (EJC core components) as well as ALYREF and DDX39B (TREX components), NCBP3 did this most efficiently; this is even more striking considering that NCBP3-LAP expression and capture levels were discernibly lower than those of NCBP1-LAP and NCBP2-LAP (Fig. 2b). NCBP3’s interaction with the EJC proteins was only mildly RNase sensitive, whereas the interaction of RBM8A and MAGOH with NCBP1 and NCBP2 were more affected by RNase treatment. It was also notable, that the EJC core component MLN51 was not identified in the NCBP3 IP-MS screening, yet it appeared in the IP-western validation (Fig. 2b, lanes 7, 9 and 11). However, compared to the three other EJC core components, MLN51 was captured less efficiently. This finding might be explained by the largely cytoplasmic localization of MLN51, whereas the remaining EJC core components and NCBP3 mainly reside in the nucleus (30,37,38). Finally, we also performed reverse-IP/western analysis using HeLa cells stably expressing HA-tagged EIF4A3 or LAP-tagged ALYREF. Both EIF4A3-HA and ALYREF-LAP IPs recapitulated previous data by co-purifying NCBP3 in an only mildly RNase-sensitive manner (Fig. 2c, 2d).

We conclude that the most abundant interactors of NCBP3 are EJC and TREX components. Unlike NCBP1 and NCBP2, NCBP3 does not copurify PHAX and exosome adaptors, suggesting that it neither functions in sn(o)RNA transport nor in exosomal RNA decay.

### EJC core integrity is required for its interaction with NCBP3

To further characterize the interaction dependence of the EJC and TREX complexes with NCBP3, we conducted NCBP3-LAP IPs upon depletion of selected EJC and TREX components. As previously reported, only partial depletion was achieved for the EJC components EIF4A3 and RBM8A (39), presumably due to the essential nature of these factors (Fig. 3a, lanes 1-3). Still, their decreased expression resulted in reduced IP levels of the remaining two EJC core factors EIF4A3 and MAGOH/RBM8A, whereas depletion of ALYREF and DDX39B had no effect (Fig. 3a, lanes 6-10). This suggests that trimeric EJC core integrity is important for NCBP3 interaction. MLN51 IP levels were unaffected by EIF4A3 or RBM8A depletion.

**Fig. 3.**
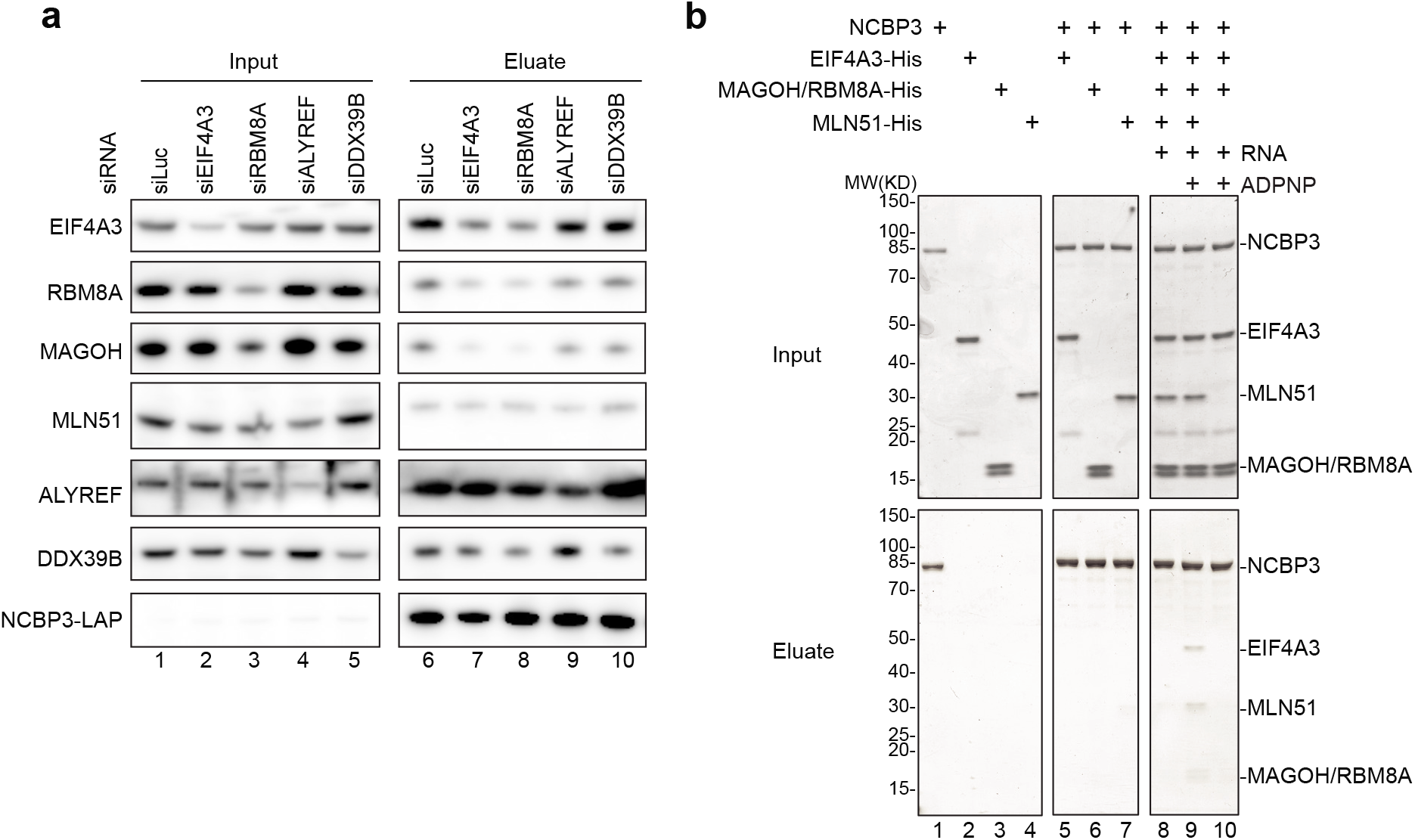
EJC core integrity is required for its interaction with NCBP3. (A) Western blotting analysis of the indicated proteins after NCBP3-LAP co-IP with anti-GFP antibody from HeLa cell extracts depleted for EJC (EIF4A3 and RBM8A) or TREX (ALYREF and DDX39B) components. Lanes 1-5 and 6-10 show input and eluate samples, respectively. Gel loading: 0.05% of input, 15% of eluate. (B) Coomassie stained SDS-PAGE showing the results of protein binding assays of NCBP3 with EJC core components added as indicated on the top. Recombinant NCBP3 was incubated with individual components or a combination of EIF4A3, MAGOH/RBM8A as a heterodimer and the selor domain of MLN51, complemented or not with ADPNP and/or single-stranded biotinylated RNA. Pull downs were then performed with the protein mix and m_7_GTP Sepharose.

We next interrogated the EJC core integrity-dependent NCBP3 interaction further, by conducting *in vitro* protein binding assays of recombinant EJC core proteins and NCBP3. Taking advantage of the affinity of NCBP3 to m_7_-GTP, we used m_7_-GTP Sepharose to purify NCBP3-bound proteins (30,32). This revealed that NCBP3 was unable to associate with either of the individual EJC core proteins (MAGOH/RBM8A added as a heterodimer) (Fig. 3b, lanes 5-7), or with a simple mix of these individual core components (Fig. 3b, lane 8). Only when the EJC core, with all four of its components, was reconstituted on RNA and locked in its closed conformation upon addition of the non-hydrolysable ATP analogue ADPNP (40), was NCBP3 able to bind (Fig. 3b, lane 9). Hence, although MLN51 interaction with NCBP3 was less abundant compared to the other EJC core components (Fig. 2b), and appeared insensitive to EIF4A3 or RBM8A depletion (Fig. 3a), the protein was still required for EJC interaction with NCBP3 *in vitro* (Fig. 3b, compare lanes 9 and 10) (see Discussion). We also performed pull down assays with an NCBP3 construct comprising only the aa1-128 N-terminal region, which harbours the RRM of the protein, conferring cap and ARS2 affinity (30,32). Although slightly less efficient than the full-length protein, this N-terminal fragment was also able to pull down the EJC core (Supplementary Fig. 3).

Taken our data together, we conclude that NCBP3 binds to the EJC, but only when it is fully assembled, which suggests that the interaction occurs post-splicing.

### NCBP3 localizes with the EJC and TREX complexes *in vivo*

Because of our established physical connection of NCBP3 to the EJC and TREX complexes, we next wondered if NCBP3 would co-localize with EJC/TREX proteins within cells. Previous immunolocalization studies showed the concentration of TREX components in splicing factor-rich nuclear speckles (41–45), and the EJC core assembling in surrounding regions, the so-called peri-speckles (38). Hence, we conducted immunolocalization analysis of NCBP3 and the nuclear speckle marker SC35, which showed a strong co-localization of the two proteins with an additional nucleoplasmic distribution of NCBP3 (Fig. 4a). A parallel comparison of co-localization of NCBP1, NCBP2 and NCBP3 with SC35, using our LAP-tagged NCBP cell lines, again showed enrichment of NCBP3 in SC35 positive nuclear speckles, whereas NCBP1 and NCBP2 distributed more evenly in the nucleoplasm (Supplementary Fig. 4). This reinforced the association of NCBP3 with splicing and RNA export activities.

**Fig. 4.**
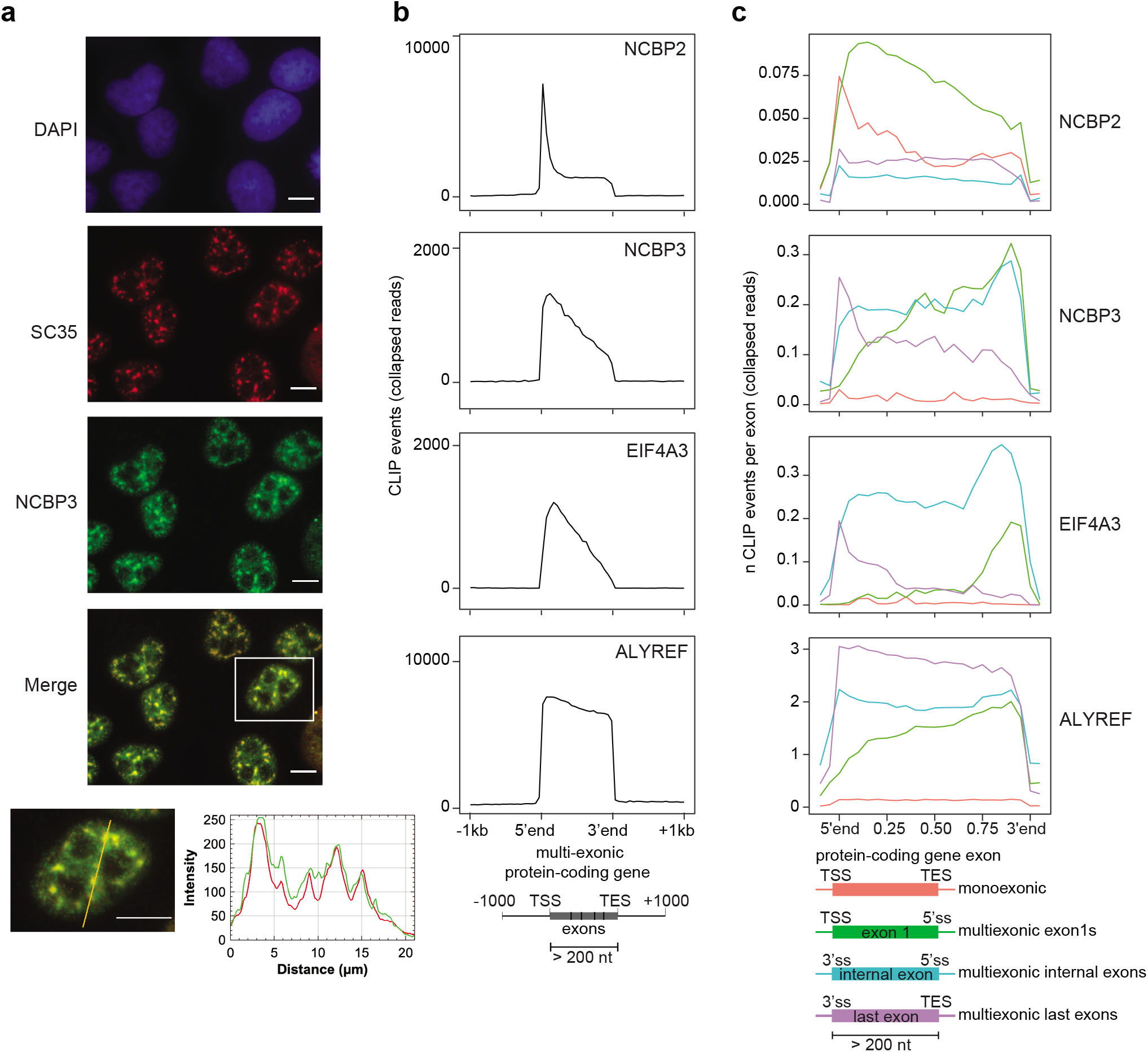
NCBP3 co-localizes with EJC and TREX complexes *in vivo*. (A) Immunolocalization microscopy of NCBP3 and the nuclear speckle marker SC35 in HeLa cells, using antibodies. DAPI, SC35 and NCBP3 stainings are shown in blue, red and green, respectively. SC35 and NCBP3 signals were merged and a zoomed-in cell is shown with a line scan profile along the drawn yellow line. Scale bars are 10 μm. (B) CLIP signal profiles of NCBP2, NCBP3, EIF4A3 and ALYREF over exons of protein coding transcripts from their transcription start sites (TSSs) to transcript end sites (TESs). (C) CLIP signal profiles as in (B) but over the exons (using a subset of exons >200nts long) of monoexonic (red) as well as first (green), internal (blue) or last (purple) exons of multiexonic protein coding transcripts.

We next investigated any preferential binding of NCBP3 to RNA by re-analyzing published CrossLinking and ImmunoPrecipitation (CLIP) data sets for NCBP3 (46,47), EIF4A3 (46,48), NCBP2 (13), and ALYREF (22). To this end, we first compared the binding profiles of these factors on exonic regions of protein coding transcripts. Unlike NCBP2, which displayed an expected cap-proximal peak, CLIP signals of NCBP3 traced further downstream into mRNA bodies, quite similar to EIF4A3 and less dramatic than ALYREF (Fig. 4b). We then examined factor-binding preferences on exons; specifically those of monoexonic- and the first, internal and last exons of multiexonic-transcripts. Expectedly, NCBP2 signal was enriched close to the caps of monoexonic and the first exons of multiexonic RNAs, whereas little signal was detected on the internal and last exons of multiexonic transcripts (Fig. 4c). In contrast, NCBP3 signal was rather low over monoexonic transcripts, whereas it was enriched close to the 3’ends of first and internal exons, and with a bias to the 5’ splice site for the last exons, much like the EJC protein EIF4A3 (Fig. 4c and Supplementary Fig. 4b). Finally, ALYREF showed an affinity, similar to NCBP3, for first exon 3’ends, but also a marked binding to last exons. Given the comparable binding profiles of NCBP3 and EIF4A3, we analysed NCBP3-RNA interaction in higher resolution, based on the thymidine to cytidine (T>C) conversions generated from protein-RNA crosslinking in the PAR-CLIP procedure. This revealed two broad peaks of NCBP3 binding on first and internal mRNA exons: one around 10-20 nt and the other 25-35 nt upstream of exon-exon junctions (Supplementary Fig. 4c). Hence, these appear just adjacent to the canonical EJC binding region, which is 20-27 nt upstream of exon-exon junctions (49–51).

We conclude that NCBP3’s physical interactions with EJC and TREX complexes are also reflected by cellular localization and RNA binding, with the RNA binding profile being particularly similar between NCBP3 and EIF4A3.

### NCBP3 depletion causes exon-number biased regulation of RNA expression

The relationship between NCBP3 and EJC/TREX prompted the question whether NCBP3 might function in the context of EJC and/or TREX. To probe a possible functional relation of NCBP3 to the EJC, we took advantage of the circumstance that depletion of EJC core components, or the peripheral component RNPS1, affects certain alternative splicing decisions (39,52). However, examining the effect of NCBP3 depletion on three exon exclusion events, that were all sensitive to EIF4A3 depletion, yielded no discernible phenotypes (Fig. 5a).

**Fig. 5.**
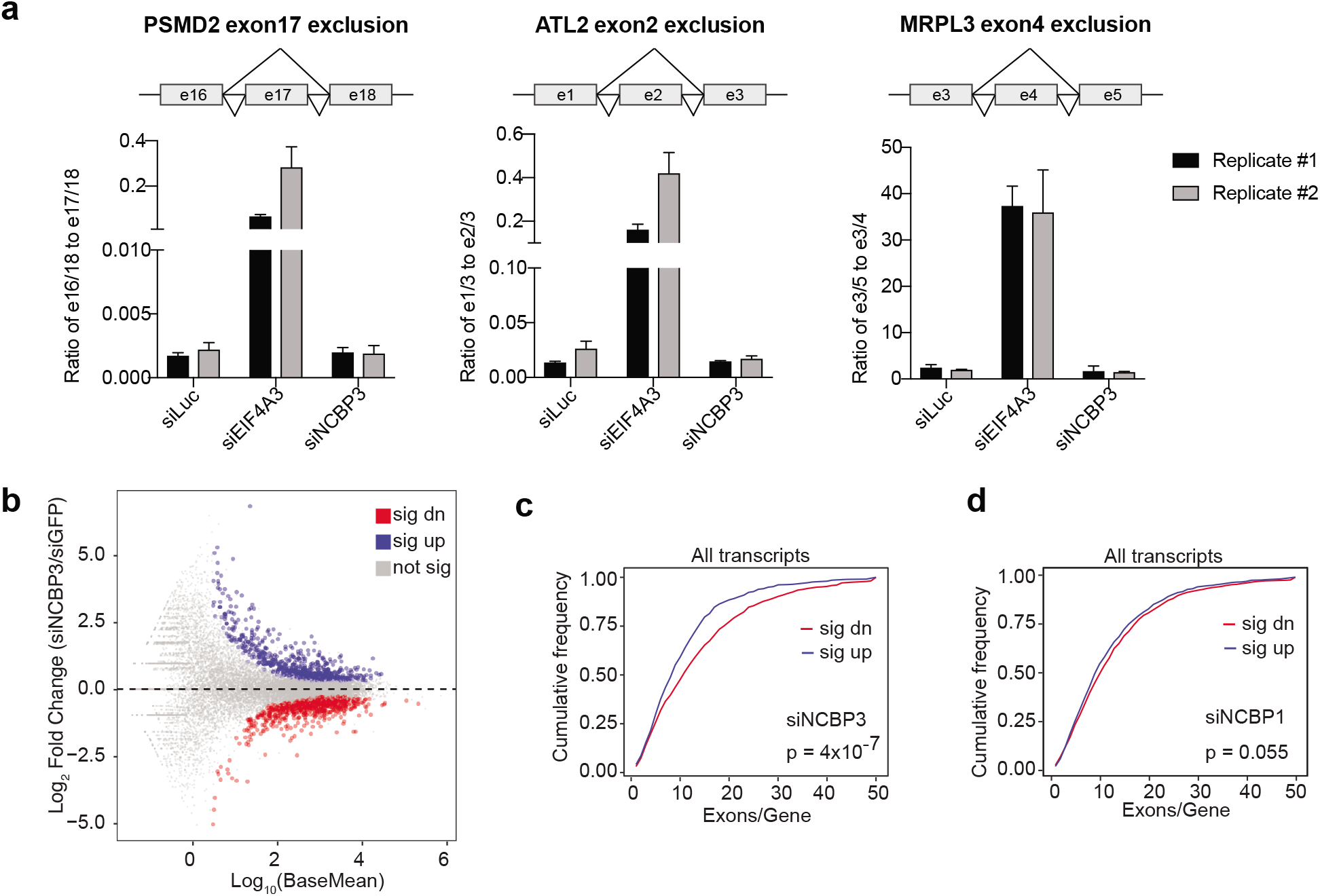
NCBP3 positively impacts the expression of multi-exonic transcripts. (A) RT-qPCR analysis of the schematized exon skipping events upon depletion of Luciferase (siLuc), EIF4A3 (siEIF4A3) or NCBP3 (siNCBP3) as indicated, and using primers listed in Supplementary Table 8. Y-axes show the ratios of exon skipping isoforms relative to non-skipping isoforms. Biological duplicate experiments are shown with error bars indicating standard deviation from technical qPCR triplicates. (B) MA-plot showing the population of significantly up- (blue) and down-regulated (red) annotated transcripts from HeLa cells subjected to NCBP3-relative to eGFP (control)-depletion. Log2 fold changes on the Y-axis are plotted against the Log10 mean of normalized counts of control and siNCBP3 samples (‘baseMean’, X-axis). Unaffected RNAs are shown in grey. (C) Cumulative frequency plot of exon numbers in upregulated (blue) and downregulated (red) transcripts upon NCBP3 depletion. The P value from a Mann-Whitney U-test comparing exon numbers in significantly up-*vs*. down-regulated transcripts is indicated. (D) Cumulative frequency as in (C) but for NCBP1 depletion.

We then asked whether NCBP3 impacts global RNA expression levels by sequencing total RNA (RNA-seq) from HeLa cells depleted of NCBP3 (Supplementary Fig. 5a and Supplementary Table 3). Differential expression analysis identified 724 and 818 annotated transcripts that were significantly up- or down-regulated upon NCBP3 depletion, respectively (Fig. 5b). We noticed that transcripts containing multiple exons (>10) were biased towards being downregulated (Fig. 5c), and that this trend was dominated by protein coding transcripts (Supplementary Fig. 5b). Employing RNA-seq data collected in parallel (53,54) demonstrated that NCBP1 depletion did not show such an exon-content effect on RNA levels (Fig. 5d, Supplementary Fig. 5b). A possible caveat to this finding was that expression levels of upregulated transcripts were significantly lower than for those that were downregulated (Supplementary Fig. 5c), and that RNAs containing many exons generally were present in higher levels than those with fewer exons (data not shown). However, using a subset of expression-matched up- and down-regulated transcripts revealed a similar trend, supporting that the phenotype of NCBP3 depletion was exon-but not expression-dependent (Supplementary Fig. 5D).

We conclude that NCBP3 does not partake in core EJC functions, yet the protein positively impacts the expression of multi-exonic transcripts. Whether this exon number-dependent effect relates to transcription, RNA decay or RNA localization remains unclear.

### NCBP3 contributes to the nuclear export of polyadenylated RNA

We then proceeded to inquire about a functional connection of NCBP3 with the TREX complex. Since individual depletion of NCBP3 does not elicit a prominent polyadenylated (PA_+_) RNA export defect (30), we sought for a possible synthetic effect between NCBP3 and TREX inactivation. Depletion of individual TREX components often results in strong nuclear accumulation of PA_+_ RNA (55), so we selected the THOC3 subunit whose depletion has a relatively mild phenotype (55), and performed fluorescent *in situ* hybridization (FISH) for PA_+_ RNA upon single- and double-depletion of NCBP3 and THOC3 (Fig. 6a and S6). Co-depletion of the TREX components DDX39A and DDX39B was conducted as a positive control (56). As demonstrated in previous studies (30,55), individual depletion of NCBP3 and THOC3 did not yield strong nuclear PA_+_ RNA accumulation (Fig. 6a). Instead, co-depletion of NCBP3 and THOC3 yielded a strong export defect. This suggests that NCBP3 plays a role in RNA export in conjunction with TREX.

**Fig. 6.**
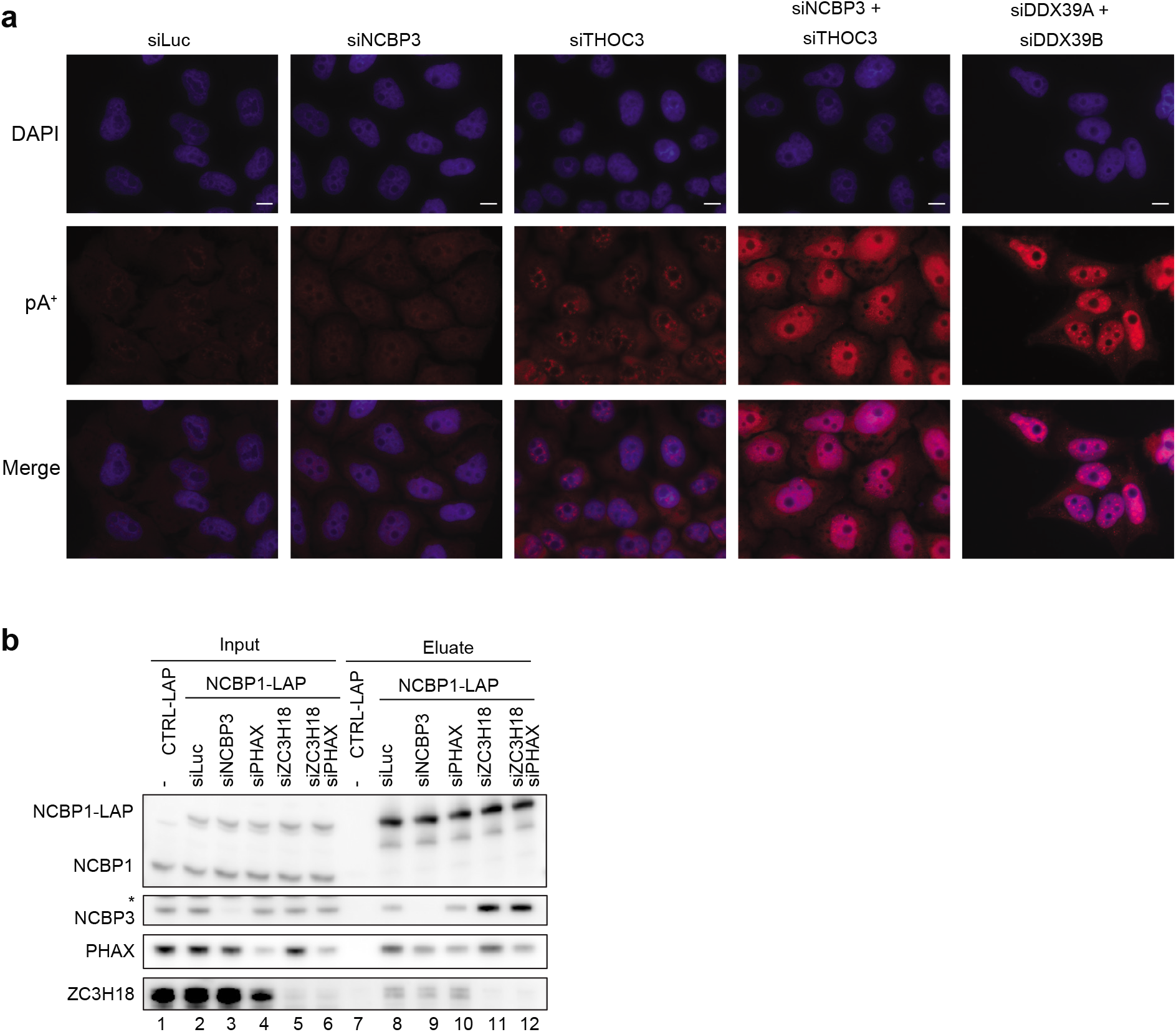
NCBP3 positively impacts PA_+_ RNP export competence. (A) Fluorescence *in situ* hybridization (FISH) analysis of PA_+_ RNA in HeLa cells after individual- or co-depletion of factors with the indicated siRNAs. Depletions were performed for 72 h with siLuc (control), siNCBP3, siTHOC3, and co-depletion with siNCBP3 and siTHOC3. For siDDX39A and siDDX39B depletion was performed for 48 h. DAPI and PA_+_ RNA signals are shown in blue and red, respectively. Merge is shown at the bottom. Scale bars are 10 μm. (B) Western blotting analysis of NCBP1-LAP co-IPs from HeLa cell extracts depleted of luciferase (control), NCBP3, PHAX, ZC3H18, or both PHAX and ZC3H18 as indicated. Lanes 1-6 and lanes 7-12 show input and eluate samples, respectively. Asterisk indicates unspecific protein band. Gel loading: 0.05% of input, 15% of eluate. IPs were performed using an anti-GFP antibody.

A function of NCBP3 in the export of PA_+_ RNA, taken together with the absence of exosome-related proteins in our NCBP3 IPs, made us suspect that NCBP3 might generally aid in the formation of ‘productive mRNP’. As PHAX and ZC3H18 were previously shown to compete for CBC binding, facilitating snRNA export and exosomal decay respectively (13), we wondered whether a similar antagonistic relationship would exist between NCBP3 and ZC3H18 for mRNA. We therefore conducted NCBP1-LAP IPs from cell extracts depleted for NCBP3, PHAX or ZC3H18. Depletion of NCBP3 did not notably affect levels of PHAX and ZC3H18 captured by NCBP1-LAP (Fig. 6b, compare lanes 8 and 9), which was probably due to the naturally lower abundance of NCBP3 associating with NCBP1 compared to that of ZC3H18 and PHAX (Fig. 2a, NCBP1 IP in condition 12 also used in the presented IPs). However, depletion of ZC3H18 led to a marked increase in levels of captured NCBP3 (Fig. 6b, lane 11), while depletion of PHAX did not change the level of captured NCBP3, and co-depletion of PHAX and ZC3H18 showed a similar effect as depleting ZC3H18 alone (Fig. 6b, lanes 10, 11 and 12). Taken together, this indicates that *in vivo* NCBP3 competes primarily with ZC3H18 for CBC binding.

## Discussion

The CBC occupies the nascent RNA cap soon after its emergence from RNAPII, which positions the complex centrally to impact RNA transcription, processing, nuclear transport and decay (5). For two decades, the CBC was known to consist of NCBP1 and NCBP2 (2). However, the previously uncharacterized protein C17orf85 was recently re-named NCBP3 due to its measurable cap-affinity and its suggested ability to form an alternative CBC dimer with NCBP1 (30). Data presented here suggest that the cap-affinity of NCBP3 is unlikely to be relevant for alternative CBC formation. Firstly, NCBP2 depletion prevents, rather than enhances, NCBP3 binding to NCBP1 (Fig. 1a). Secondly, NCBP3 does not bind RNA in a cap-proximal manner expected by a CBC component and as displayed by NCBP2 (Fig. 4c). Thirdly, NCBP3 associates with both NCBP1 and NCBP2, albeit only weakly (Fig. 2a), as reported previously (30,32), making it more likely that NCBP3 interacts with the canonical CBC *in vivo*. Consistently, our interaction profiling of the three NCBPs demonstrated that NCBP3 engages in interaction networks distinct from those of NCBP1 and NCBP2, and instead pointed to the EJC and TREX complexes as its most dominant interactors. NCBP3 localization and functional analyses further strengthened this connection. Based on these data, we therefore propose a model for the involvement of NCBP3 in nuclear RNA metabolism (Fig. 7). The CBC-bound RNA is receptive to the binding of different productive and destructive factors, which dynamically exchange during RNP formation to ultimately define transcript fate (28). PHAX and ZC3H18 are established examples of such RNP components, which compete for CBC-binding and ultimately tilt RNA fate towards snRNP/snoRNP assembly or nuclear degradation by the exosome, respectively (13). Based on its abilities to compete with ZC3H18 for NCBP1 binding and to stimulate PA_+_ RNA nuclear export (Fig. 6), we suggest that NCBP3 is involved in such nuclear RNP definition, favouring EJC-bound transcripts to become export competent.

**Fig. 7.**
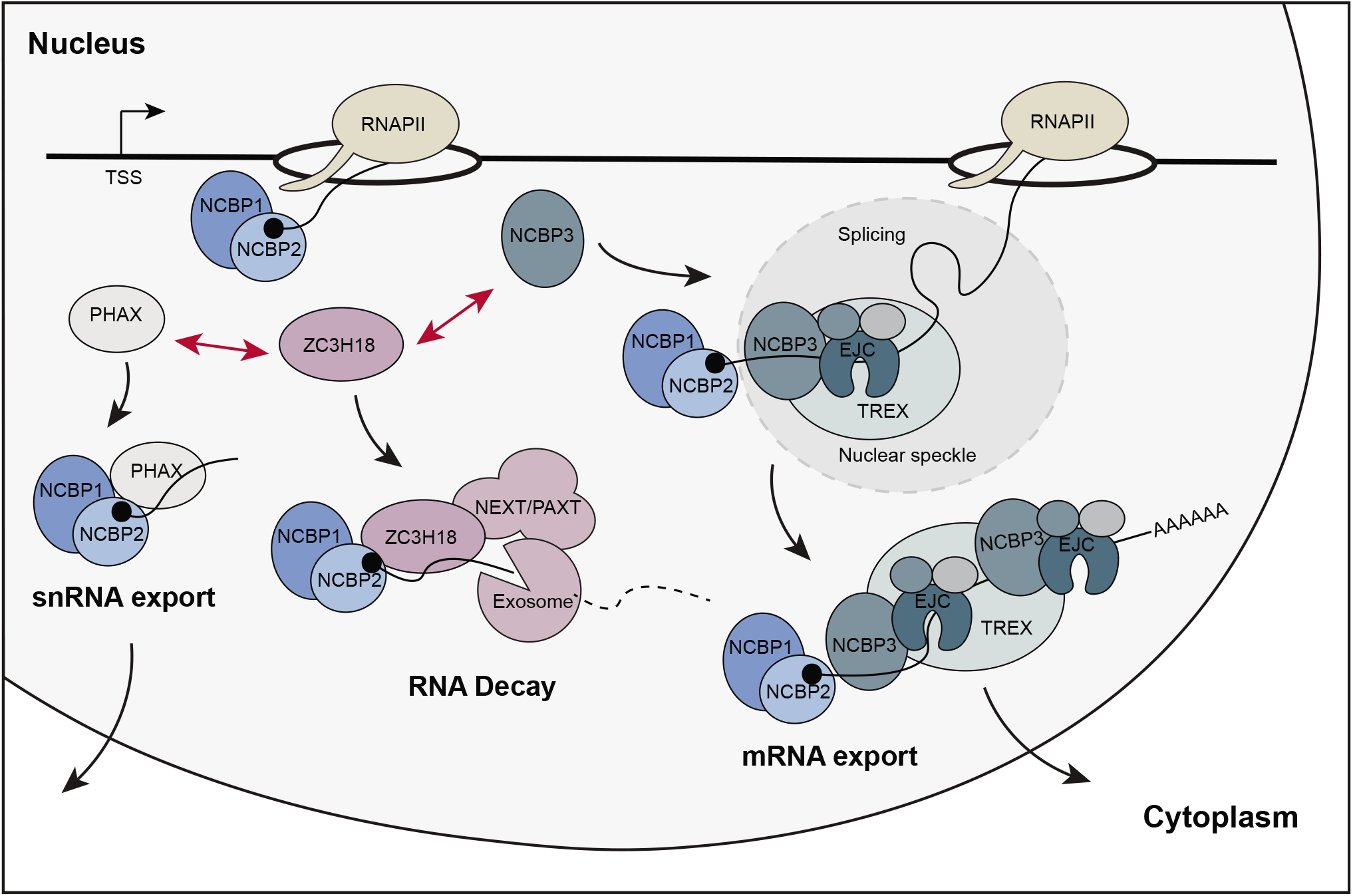
Model for NCBP3 involvement in RNA fate decisions. Shortly after transcription initiation the CBC (NCBP1 and NCBP2) binds the RNA cap. Hereafter, the nascent RNA is exposed to different factors, which compete for RNA binding (exemplified by PHAX, ZC3H18 and NCBP3). As shown in the present paper, NCBP3 competes with ZC3H18 for CBC interaction, which is likely tilted in the favour of NCBP3 by its association with EJC and TREX. As such, NCBP3 contributes to licensing PA_+_ RNPs for nuclear export.

Our protein-protein interaction profiles provided a direct comparison of the binding partners of the three NCBPs. In contrast to the comparable NCBP1 and NCBP2 interaction profiles, the NCBP3 interaction screening only detected NCBP1 and NCBP2 in low amounts and neither picked up PHAX nor the exosome-related factors ZC3H18, ZCCHC8 and MTR4 (Fig. 2a). Instead, NCBP3 abundantly co-purified constituents of the EJC. Curiously though, while most of the EJC and its peripheral components were detected, the EJC core component MLN51 was absent. As an NCBP3-MLN51 interaction was independently established by IP/western blotting analysis (Fig. 2b), it is possible that a technical issue might explain the lack of an MLN51 MS signal, which was also suspiciously absent in previous mRNP IPs (57,58). However, it could also reflect a relatively low abundant MLN51-NCBP3 interaction. On this note, it was suggested that not all of the trimeric EJC (eIF4A3, MAGOH, RBM8A) associates with MLN51, since the protein was substoichiometric to the other EJC core components in both EIF4A3 and MAGOH IPs (59). Moreover, a recent study demonstrated that RNPS1, which was also identified in our NCBP3 interaction screen, and MLN51 associate with the trimeric EJC in a mutually exclusive manner, forming early (mainly nuclear) and late (mainly cytoplasmic) EJCs, respectively (60). This might also help explain why MLN51 is required for NCBP3-EJC interaction *in vitro*, which is not obvious *in vivo*, as the reconstituted environment lacking other proteins, such as RNPS1, that could influence EJC formation. Interactions with MLN51 notwithstanding, the association of the bulk of NCBP3 with EJC was also reflected by their similar RNA binding profiles derived from CLIP datasets; NCBP3 binds close to the splice junctions of multiexonic RNAs, showing two broad peaks at 10-20 nt and 25-35 nt upstream of exon-exon junctions (Fig. 4c and Supplementary Fig. 4c). This would be consistent with binding around a deposited EJC, which covers around 6-9 nt and a prominent EIF4A3 cross-linking site at ~27 nt upstream of the splice junction (40,50,51,61). Based on this, we speculate that the EJC may direct NCBP3 binding (see below).

Consistent with its physical interaction with EJC proteins, we found NCBP3 co-localizing with nuclear speckles (Fig. 4a), which are also enriched for pre-mRNA splicing factors (45). A previous study suggested that pre-mRNA splicing is required for the association of NCBP3 with mRNA *in vitro*, as NCBP3 was absent from intron-depleted mRNPs (58). Further linking NCBP3 to splicing was the establishment of a peripheral association of the protein with the spliceosomal B and C complexes (62). However, notwithstanding these indications, we were not able to establish a role for NCBP3 in alternative splicing, using an EJC-sensitive assay (Fig. 5a). We therefore surmise that NCBP3’s interaction with the EJC does not reflect an EJC-like function of the protein. Consistent with this idea, although loading of NCBP3 onto mRNA was deemed intron-dependent, it did not require the presence of the EJC for mRNP association (58). Here, we found that NCBP3 requires a fully assembled EJC for *in vitro* interaction (Fig. 3b), and that trimeric EJC core integrity is critical for the interaction *in vivo* (Fig. 3a). Taking these results together allows us to speculate about the timing of NCBP3 recruitment to the early RNP and its subsequent interaction with the EJC. Trimeric EJC core assembly on RNA occurs after the first step of splicing before exon-exon ligation (63). The EJC-independent RNA binding of NCBP3 (58), would therefore occur between the onset of spliceosome assembly (before EJC recruitment) and until after the first step of splicing (simultaneous with EJC recruitment). Which exact molecular interactions would drive this initial RNA-NCBP3 association remains to be investigated. However, as mentioned above, we suggest that stable EJC formation ultimately provides an anchor point for NCBP3.

In addition to the EJC, NCBP3 also associates robustly with most known components of the TREX complex, including the complete THO complex (Fig. 2a). Previously, NCBP3 was copurified with TREX components ALYREF, DDX39B, THOC2 and CIP29, and it was even considered a putative new TREX subunit (29,41). However, no functional consequence of its association with TREX was reported. Here, we provide evidence for a role of NCBP3 in PA_+_ RNA nuclear export by acting redundantly with the TREX component THOC3. Still, we do not consider NCBP3 a central TREX component as its independent depletion does not elicit nuclear accumulation of PA_+_ RNA. Instead, NCBP3 may operate tangentially to TREX and this relationship may only be revealed when cells are subjected to the appropriate double depletions of factors. Perhaps the similar phenotype observed upon dual NCBP2 and NCBP3 depletion (30) reflects a related synthetic effect between co-operating RNP factors? In further agreement with an involvement of NCBP3 in the nuclear export of at least some transcripts, we found that NCBP3 positively impacts the expression of a subset of genes. Specifically, upon NCBP3 depletion, mRNAs deriving from multi-exonic transcripts were prone to be downregulated (Fig. 5c). We speculate that an abundant number of introns might increase RNA nuclear residence time and therefore require NCBP3 to fend off nuclear decay activities (as exemplified by its competition with ZC3H18), while providing additional export capacity. As shown previously in both yeast and human cells, efficient nuclear export is required to evade nuclear decay by the RNA exosome (29,64,65). Loss of NCBP3 might therefore de-stabilize these multi-exonic transcripts dually due to their slowed nuclear export and enhanced turnover (Fig. 7).

If NCBP3 is not a *bona fide* CBC protein, what is then the functional significance of the m_7_G cap-affinity of the protein? It may bind the cap when CBC is not available at certain undetermined circumstances. Alternatively, and in line with its internal mRNA association, it might potentially contact m_7_G modifications, which are found in mRNAs (66,67), but not in snRNAs and snoRNAs (68). Regardless and setting the cap affinity of NCBP3 aside, we suggest NCBP3 partakes in dynamic nuclear RNP metabolism, promoting the productive fate of mRNP.

## Data Availability

The MS proteomics data have been deposited to the ProteomeXchange Consortium via the PRIDE partner repository with the dataset identifier PXD016038. RNA-seq data have been deposited to the Gene Expression Omnibus (GEO) under the accession number GSE99059.

## Acknowledgements

We thank the National Center for Dynamic Interactome Research for facility support for the interaction screen, Dr. Ina Poser and Prof. Anthony A. Hyman for providing the LAP-tagged cell lines, and Dr. Catherine-Laure Tomasetto for providing the MLN51 antibody.

## Author contributions

Y.D., T.H.J., J.L. and H.L.H. designed the experiments. Y.D., I.B., H.J., C.I., K.R.M., W.M.S. performed the experiments. Y.D., M.S. and K.R.M. performed data analysis.

M.S., H.L.H., and S.C. provided critical input. T.H.J. and J.L. supervised the project. Y.D. and T.H.J. wrote the manuscript with input from all co-authors.

