## Supplementary for "NCBP3 is a productive mRNP component"

**Supplementary Fig. 1**

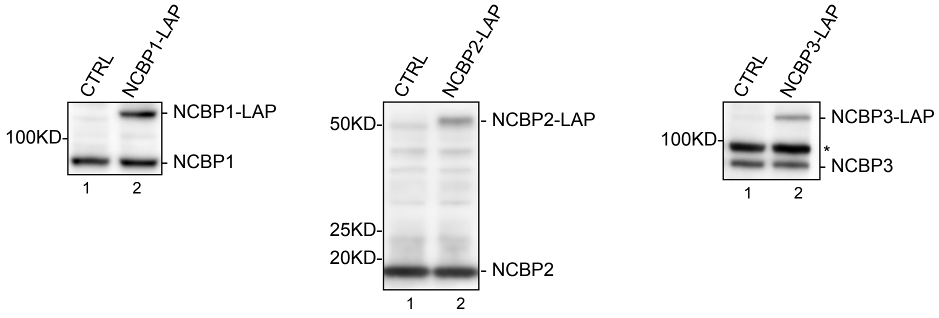


**Supplementary Fig. 1 Related to Fig. 1.**
Western blotting analysis of both endogenous and LAP-tagged NCBP proteins expressed in HeLa cells, using antibodies targeting NCBP1 (left), NCBP2 (mid) or NCBP3 (right). For each blot, lane 1 shows a non-tagged NCBP control. Asterisk indicates unspecific protein band.

### Supplementary Fig. 2

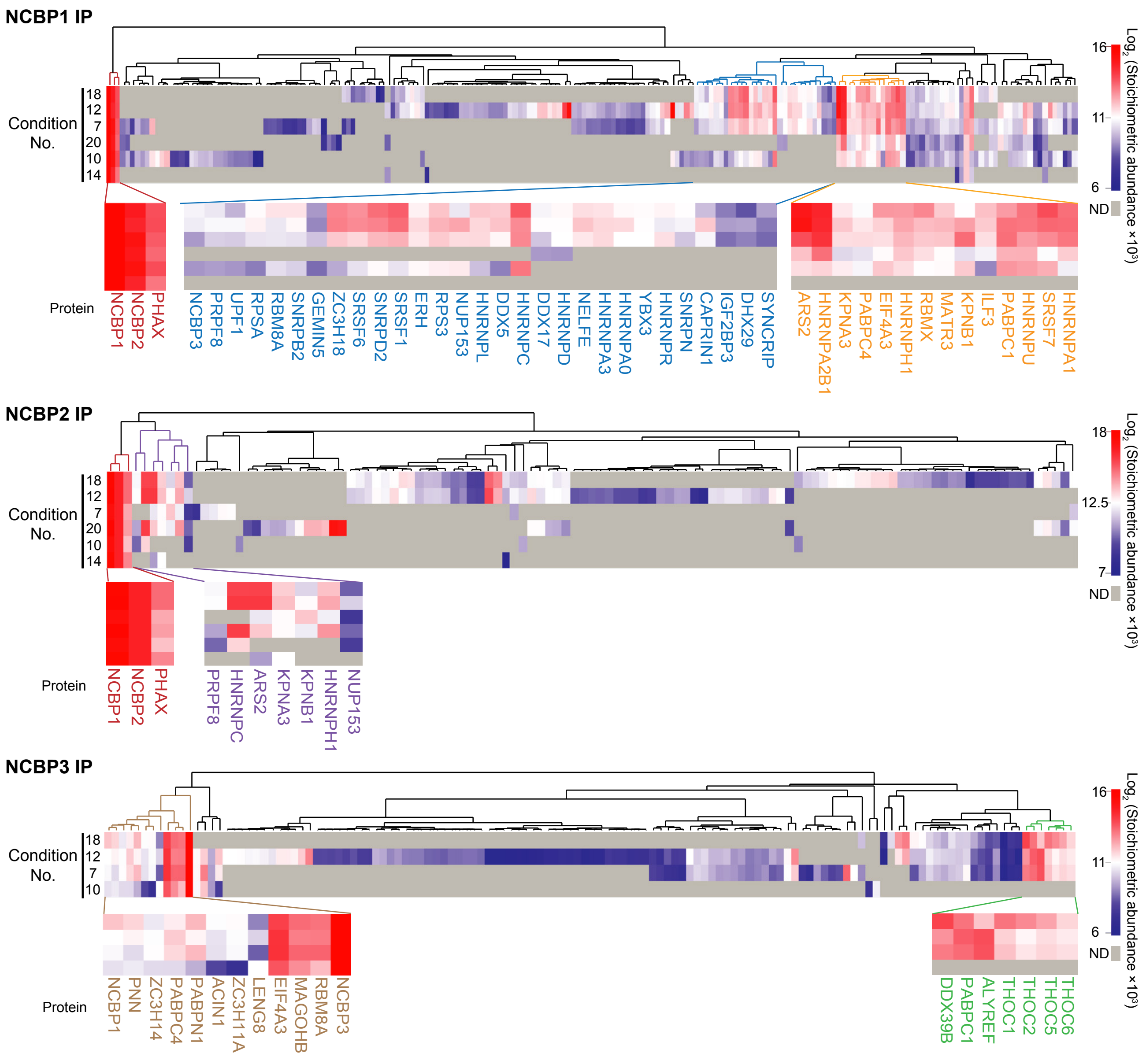


**Supplementary Fig. 2 Related to Fig. 2.**
Heatmap clustering of Euclidean distances showing log2 transformed stoichiometric abundances of NCBP1 (top), NCBP2 (mid) and NCBP3 (bottom) IPs. Missing values (ND) were imputed with 0 for the clustering. The heat colour range is shown on the right. Clusters where most proteins have abundance higher than the midpoint (in the red to white colour range) are highlighted, and zoomed-in versions with gene names are shown below the full heatmaps.

#### Supplementary Fig. 3

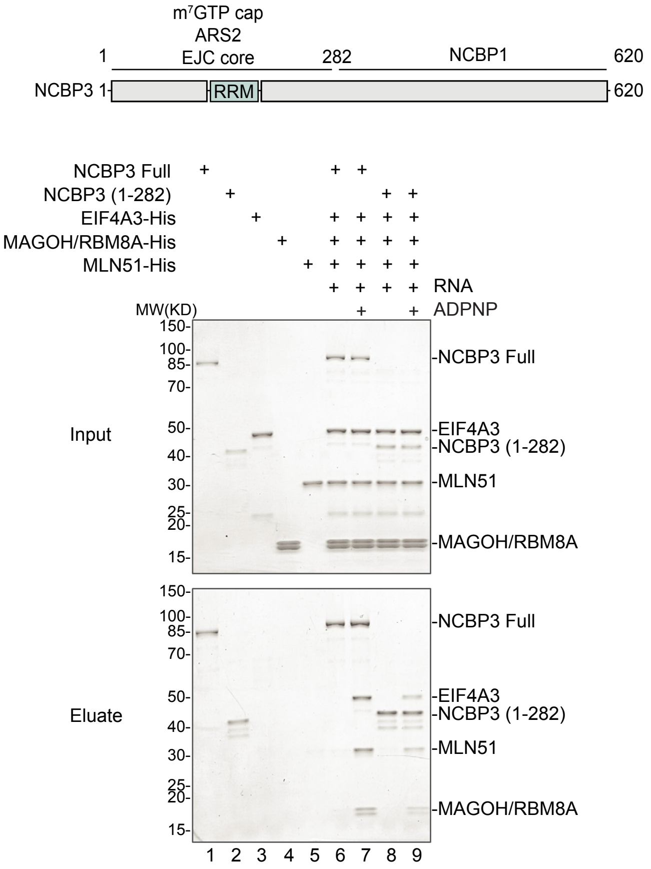


**Supplementary Fig. 3 Related to Fig. 3.**

Coomassie stained SDS-PAGE showing the results of protein binding assays of NCBP3 with EJC core components added as indicated on the top. Recombinant full length or N-terminal (1-128) NCBP3 (see schematics on the top) was incubated with EJC core components EIF4A3, MAGOH/RBM8A as a heterodimer and the selor domain of MLN51, and complemented with single-stranded biotinylated RNA with/without ADPNP. Pull-down assay were performed with the protein mix and m7GTP Sepharose.

#### Supplementary Fig. 4

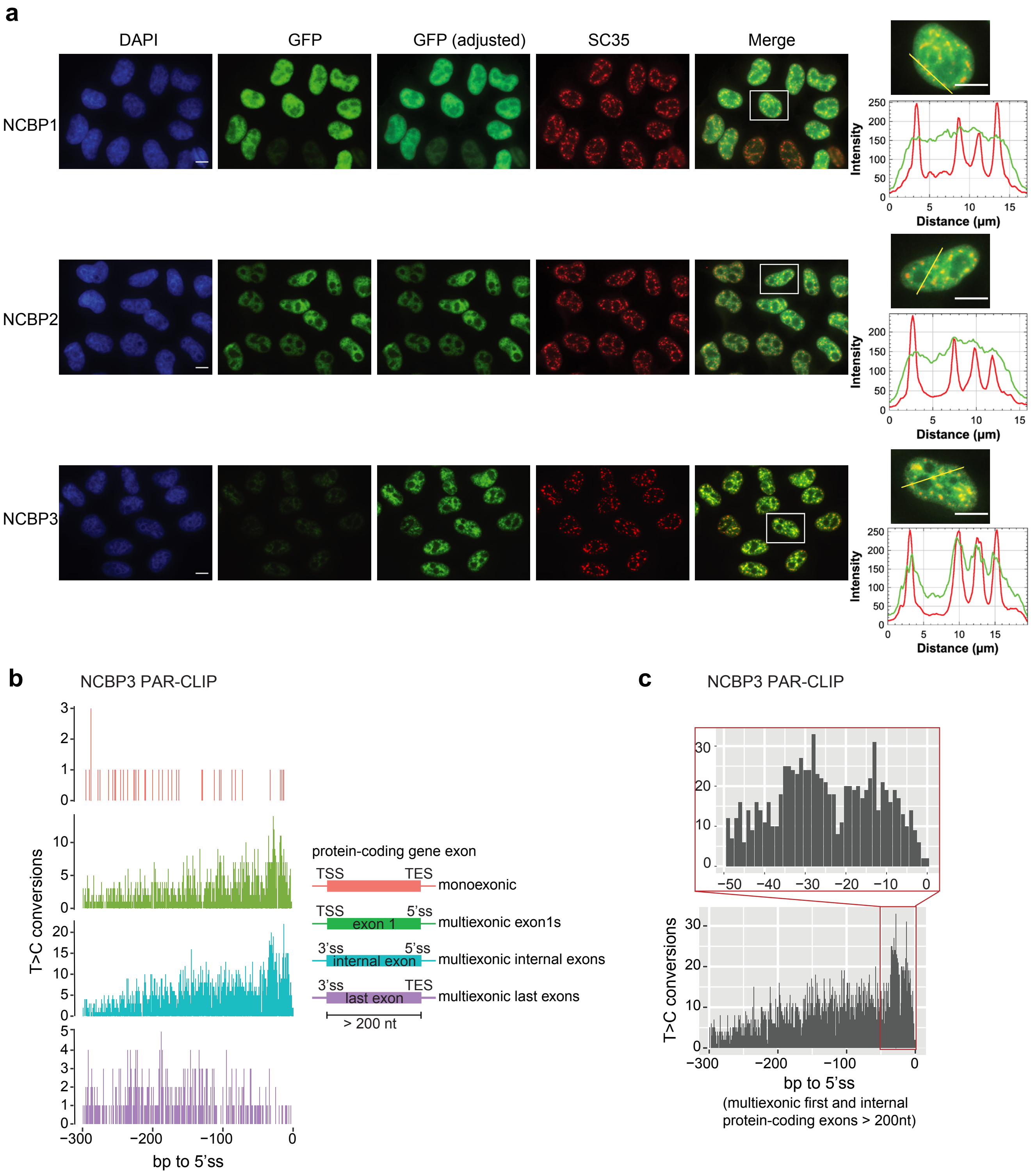


**Supplementary Fig. 4 Related to Fig. 4.**
(a) Immunolocalization microscopy of LAP-tagged NCBP1, NCBP2 and NCBP3 with SC35, using antibodies targeting GFP and SC35. DAPI, SC35 and NCBP3 signals are displayed as in Fig. 4a. GFP brightness/contrast was set similarly across all NCBPs (second column) and also adjusted to achieve better visualization of individual NCBPs (third column). Merges were performed between adjusted GFP- and SC35-signals and signal scans were performed as in Fig. 4a. Scale bars, 10 µm.
(b) NCBP3 PAR-CLIP T>C conversions on monoexonic exons as well as first, internal and last exons of multiexonic transcripts.
(c) A zoomed-in version of NCBP3 PAR-CLIP T>C conversions over multiexonic first and internal protein coding exons.

#### Supplementary Fig. 5

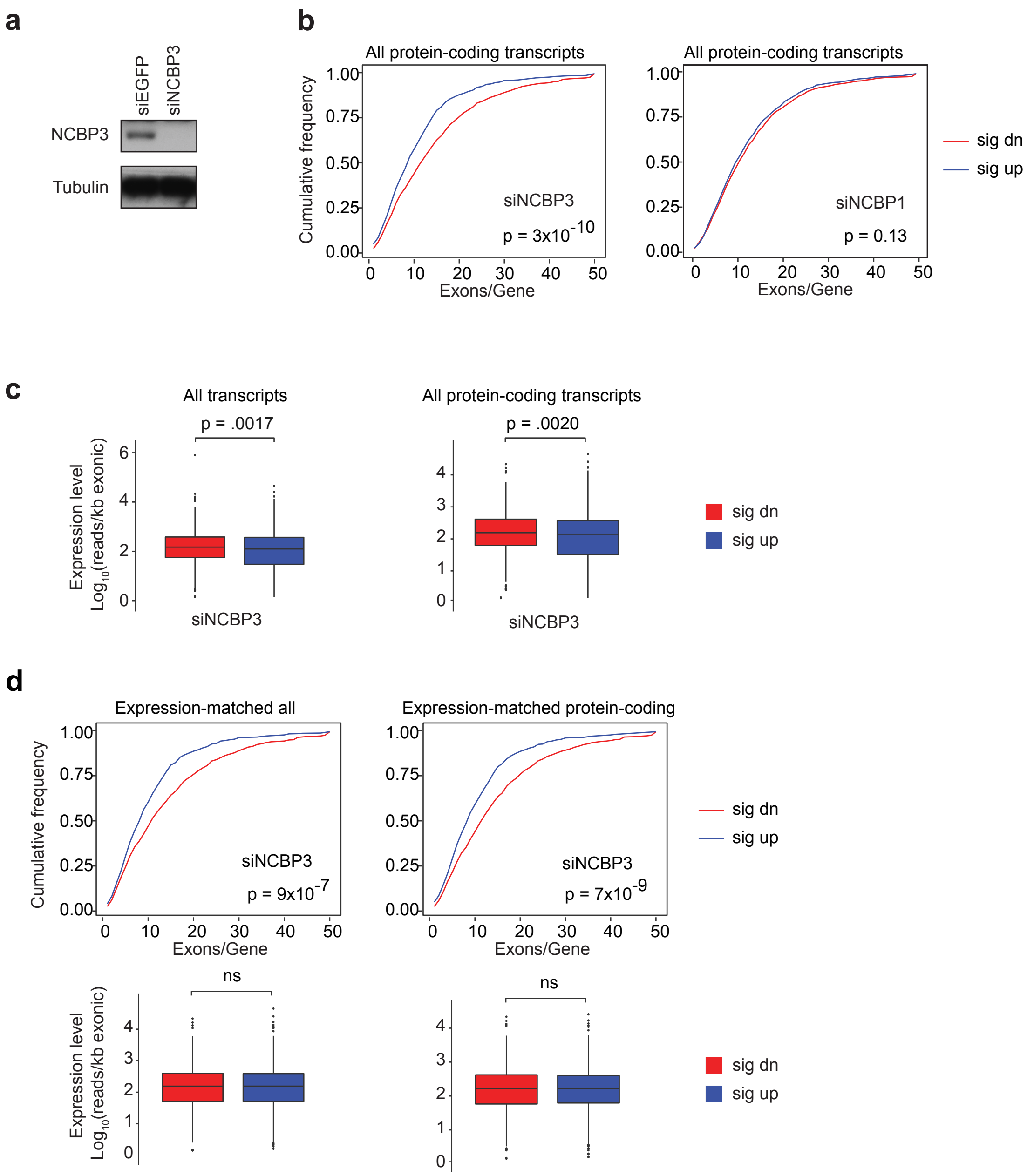


**Supplementary Fig. 5 Related to Fig. 5.**
(a) Western blotting analysis showing the depletion efficiency of NCBP3.
(b) Cumulative frequency plot of exon numbers in upregulated (blue) and downregulated (red) transcripts upon NCBP3 (left) and NCBP1 (right) depletion, as in Fig. 5c and 5d, but for all differentially expressed protein coding transcripts.
(c) Expression level comparison between significantly up- and down-regulated transcripts. Expression levels per transcript were computed as mean read coverages per kb of exonic parts of transcripts for significantly up- (blue) and down-regulated (red) transcripts as in Fig. 5c. The P value from a Mann-Whitney U-test comparing expression between significantly up vs. downregulated transcripts is indicated.
(d) Analysis of the correlation between exon numbers and changed expression upon NCBP3 depletion as in Fig. 5c, but for an equal-sized subset of expression-matched transcripts (n=675, on the left panel) and expression-matched protein coding transcripts (n=603, on the right panel). The cumulative frequency plots and p values are obtained as described in Fig. 5c.

#### Supplementary Fig. 6

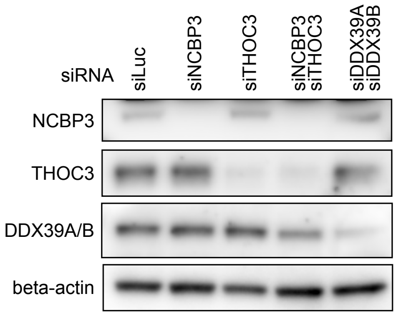


**Supplementary Fig. 6 Related to Fig. 6.**
Western blotting analysis showing protein depletion efficiencies with indicated siRNAs.

**Supplementary Table 1 Composition of extraction solutions used for the interaction screen. Related to Fig. 1 and 2, and experimental procedures.**

**Supplementary Table 2 Stoichiometric abundance of proteins enriched with NCBP1, 2 and 3 in the IP screen. Related to Fig. 1, Fig. 2 and Supplementary Fig. 2. Detailed description of the table is included in the spreadsheet.**

**Supplementary Table 3 Statistics of the RNA-seq library. Related to Fig. 5.**

**Supplementary Table 4 BACs used for the cell lines. Related to experimental procedures.**

**Supplementary Table 5 List of siRNAs used. Related to experimental procedures.**

**Supplementary Table 6 Antibodies used for Western blots. Related to experimental procedures.**

**Supplementary Table 7 Antibodies used for immunofluoresence. Related to experimental procedures.**

**Supplementary Table 8 Primers used for qPCR. Related to experimental procedures.**

**Supplementary Table 1 Composition of extraction solutions used for the interaction screen. Related to Fig. 1 and 2, and experimental procedures.**

| Condition No. | Composition |
|---|---|
| 7 | 20 mM HEPES-Na, pH 7.4, 150 mM NaCl, 5mM d-BCHAP |
| 10 | 20 mM HEPES-Na, pH 7.4, 300 mM NaCl, 0.5mM Sarcosyl |
| 12 | 20 mM HEPES-Na, pH 7.4, 150 mM NaCl, 0.5% TRIT X100 |
| 14 | 20 mM HEPES-Na, pH 7.4, 50 mM MgCl2, 0.5% TRIT X100 |
| 18 | 125 mM NH4 Acet, pH 7, 100 mM NaCl, 1% TRIT X100 |
| 20 | 20 mM TRIS-Cl, pH 8, 125 mM Na3 Cit, 1% TRIT X100 |

**Supplementary Table 3 Statistics of the RNA-seq library. Related to Fig. 5.**

| Library | Total reads | Trimming and read quality control | | Mapping | |
|---|---|---|---|---|---|
| | | read pairs remaining | | uniquely aligned, proper read pairs | |
| siEGFP_1 | 41332086 | 38139659 | (92.28%) | 29984813 | (78.62%) |
| siEGFP_2 | 52427151 | 48776569 | (93.04%) | 37667679 | (77.22%) |
| siEGFP_3 | 40868991 | 37865523 | (92.65%) | 23137272 | (61.10%) |
| siNCBP1_1 | 37239952 | 34934956 | (93.81%) | 27799828 | (79.58%) |
| siNCBP1_2 | 41266225 | 38296661 | (92.80%) | 24757893 | (64.65%) |
| siNCBP1_3 | 35493503 | 33126807 | (93.33%) | 25686591 | (77.54%) |
| siNCBP3_1 | 45221352 | 42090840 | (93.08%) | 31615218 | (75.11%) |
| siNCBP3_2 | 34333229 | 31658743 | (92.21%) | 21747355 | (68.69%) |
| siNCBP3_3 | 45619227 | 42397189 | (92.94%) | 32942139 | (77.70%) |

**Supplementary Table 4 BACs used for the cell lines. Related to experimental procedures.**

| Cell line | Gene Name | BAC ID | Tagging cassette | Ensembl ID |
|---|---|---|---|---|
| CTRL-LAP | - | CTD-3000G10 | N-terminal | - |
| NCBP1-LAP | NCBP1 | RP11-17A21 | C-terminal | ENSG00000136937 |
| NCBP2-LAP | NCBP2 | HS.E139.A23 | C-terminal | ENSG00000114503 |
| NCBP3-LAP | NCBP3 | RP11-118B14 | C-terminal | ENSG00000074356 |
| ALYREF-LAP | ALYREF | RP11-634L10 | C-terminal | ENSG00000183684 |

**Supplementary Table 5 List of siRNAs used. Related to experimental procedures.**

| Target | Sequence (sense 5'-3') |
|---|---|
| CTRL (Luciferase) | CUUACGCUGAGUACUUCGAdTdT |
| NCBP1 | GGAAAGGAGUUGUACGAAAdTdT |
| NCBP2 | GGCCAGAACUUGAGUAUUUdTdT |
| NCBP3 | UCAGCGGGACGUGAUCAAGAAdTdT |
| DDX39A | AAAGGCCUAGCCAUCACUUUUdTdT |
| DDX39B | AAGGGCUUGGCUAUCACAUUUdTdT |
| ALYREF | UGGGAAACUGCUGGUGUCCAAdTdT |
| EIF4A3 | AGACAUGACUAAAGUG GAAdTdT |
| RBM8A/Y14 | CGCUCUGUUGAAGGCUGGAdTdT |
| THOC3 | GUGUGAGUCUCCGACCUUCdTdT |

**Supplementary Table 6 Antibodies used for Western blots. Related to experimental procedures.**

| Target | Origin | Source | Cat #/reference | Working dilution |
|---|---|---|---|---|
| NCBP1 | Rabbit | E.Izzauralde | | 1:1000 |
| NCBP2 | Rabbit | E.Izzauralde | | 1:1000 |
| NCBP3 | Rabbit | Sigma | HPA013195 | 1:1000 |
| GFP (B-2) | Mouse | Santa Cruz | sc-9996 | 1:1000 |
| ALYREF | Rabbit | Abcam | Ab202894 | 1:10000 |
| EIF4A3 | Rabbit | H.Le Hir | | 1:1000 |
| RBM8A/Y14 | Rabbit | H.Le Hir | | 1:500 |
| MLN51 | Rabbit | C.L.Tomasetto | Degot et al., 2002 | 1:1000 |
| MAGOH | Mouse | Santa Cruz | sc-56724 | 1:500 |
| DDX39A/DDX39B | Mouse | Santa Cruz | sc-271395 | 1:500 |
| HA | Rabbit | Abcam | ab9110 | 1:5000 |
| THOC3 | Rabbit | Sigma | HPA044009 | 1:500 |

**Supplementary Table 7 Antibodies used for immunofluoresence. Related to experimental procedures.**

| Target | Origin | Source | Cat #/reference | Working dilution |
|---|---|---|---|---|
| NCBP3 | Rabbit | Atlas Antibodies | HPA013195 | 1:500 |
| SC35 | Mouse | Abcam | ab11826 | 1:1000 |
| GFP | Rabbit | Invitrogen | A-6455 | 1:1000 |
| Mouse (Alexa Fluor 488) | Goat | Invitrogen | A11034 | 1:1000 |
| Rabbit (Alexa Fluor 647) | Goat | Invitrogen | A21244 | 1:1000 |
| Mouse, (Alexa Fluor 633) | Goat | Invitrogen | A21052 | 1:1000 |
| Rabbit (Alexa Fluor 488) | Donkey | Invitrogen | A21206 | 1:1000 |

**Supplementary Table 8 Primers used for qPCR. Related to experimental procedures.**

| Primer name | Sequence (5'-3') |
|---|---|
| PSMD2 Exon16_18.fw | GCACGTAATTCACACAACTGAC |
| PSMD2 Exon17.fw | CAGCTCCTGCCACTCCTTAG |
| PSMD2 Exon18.rev | GCTGCTGACATCATCTCCGT |
| ALT2 Exon1_3.fw | CGACCTCCCTAGGATTCTCA |
| ALT2 Exon2.fw | GTAGTGGTATCTGTGGCAGGAG |
| ALT2 Exon3.rev | CTCGCCATGTAAAGCCTGTCA |
| MRPL3 Exon3_4.fw | CATTACTTCAGGTACAAGAC |
| MRPL3 Exon3_5.fw | CATTACTTCAGAAAGCTACATCC |
| MRPL3_Exon5.rev | GCCTGGTTTAATTGCAGCAT |
